# Microclimate and radiant energy measurements using a novel device for natural resource monitoring

**DOI:** 10.1101/072108

**Authors:** Taylor Thomas, Adam Wolf

## Abstract

A novel device for microclimate and radiant energy monitoring of natural resources was compared to known reference instruments and sensors to gauge relative performance under field conditions in Nebraska and New Jersey, USA during summer 2016. For all measurements tested, including air temperature, relative humidity, barometric pressure, four component net radiation, and photosynthetically active radiation, the device reported values near or within the accuracy limits as stated by the reference instrument / sensor manufacturer.

## INTRODUCTION

Plants live within the bounds of a physical environment that is primarily controlled by local atmospheric conditions, radiant energy, and water and nutrient availability (2). The accurate measurement of ambient air temperature (*T*_*a*_), relative humidity (*RH*), and barometric pressure (*P*_*a*_) above the Earth’s surface provides insights into the dynamics of near-surface meteorology and the plant status in the ground-plant-atmosphere continuum of matter and energy. The development of heat stress (1), development of water stress (3), and plant productivity (2) are functions of processes that are controlled by these parameters. The radiation budget at a point can be said to be composed of radiation originating from the sun, termed shortwave radiation, and radiation from the organic and inorganic components of the Earth system, termed longwave radiation. Net radiation, or total incoming subtracting total outgoing radiation, is the quantity of energy available to drive natural processes such as sensible heat exchange, latent heat exchange, and photosynthesis (10). Photosynthetically active radiation (PAR) is the total incident radiation within the 400 nm to 700 nm waveband. Radiation, typically expressed as photon flux density (4) within this waveband is available to plants to drive photosynthetic machinery, the process by which carbon dioxide is assimilated into plant biomass (7). Tables 1 - 2 shows a summary of the accuracy of device measurements, relative to references.

## MATERIALS AND METHODS

### Air Temperature

The Pulsepod measures *T*_*a*_ within a carefully designed chamber that houses ambient air temperature sensors, including a thermistor and a band-gap temperature sensor, within the air stream below the device. This design serves to thermally isolate the device body from the measurement location, ensuring that the measured *T*_*a*_ is representative of the bulk air surrounding the device. The chamber vents allow air to flow freely over the sensor so that stagnant air (10) is not surrounding the temperature sensor. A Pulsepod measured *T*_*a*_ over a soybean canopy (*Glycine max*) alongside a reference Vaisala HMP50 Temperature and Relative Humidity Sensor (*Vaisala Instruments, Vantaa, Finland*) housed within a radiation shield and mounted at the same measurement height over the canopy (Fig. 1 - 2). The Vaisala HMP50 temperature sensor is a 1000 Ω platinum resistance thermometer with a manufacturer stated accuracy of ± 1.5°C across the temperature range -40°C to 60°C (12). Data shown were collected at the University of Nebraska Carbon Sequestration Program (b) Site 3 at Mead, NE in June, 2016. Device measurement height was five meters. Site coordinates are: 41.179665, -96.439336.

### Relative Humidity

Relative humidity (*RH*) is defined as “the ratio of actual amount of water vapor in the air compared to the equilibrium (saturation) amount at a given temperature” (8). *RH* is in part a function of the ambient air temperature, so the accuracy of the temperature measurement propagates into the accuracy of the *RH* measurement. The Pulsepod measures relative humidity in the same chamber as it measures air temperature, using a capacitive detection method. The location within the chamber permits ongoing *RH* measurements of free air around the device body. The reference *RH* sensor is the Vaisala INTERCAP capacitive chip (12). Data shown were collected at the University of Nebraska Carbon Sequestration Program (b) Site 3 at Mead, NE in June, 2016 (same as air temperature reference location).

### Barometric Pressure

Barometric pressure (*Pa*), or atmospheric pressure, is defined as “the total force per unit area acting isotropically on a body within the atmosphere” (11). It is the resultant force of all atmospheric particles above a sensor location. With an increase in elevation, the total atmospheric pressure is reduced, while the short-term fluctuations in *Pa* are closely tied to changing weather conditions. The Pulsepod measures *Pa* with a compact piezoresistive sensor that responds to changes in local pressure conditions. The *Pa* reported by the Pulsepod is compared to the *Pa* as reported by the reference sensor, RM Young 61302V (*RM Young, Traverse City, MI*) with a manufacturer stated accuracy of 0.05% (± 1 σ) over the analog pressure range (9). Data shown were collected at the Rutgers University Photochemical Assessment Monitoring Site (a) at New Brunswick, NJ in August, 2016. Device measurement height was two meters. Site coordinates are: 40.462204, -74.429499.

**Figure 1.**
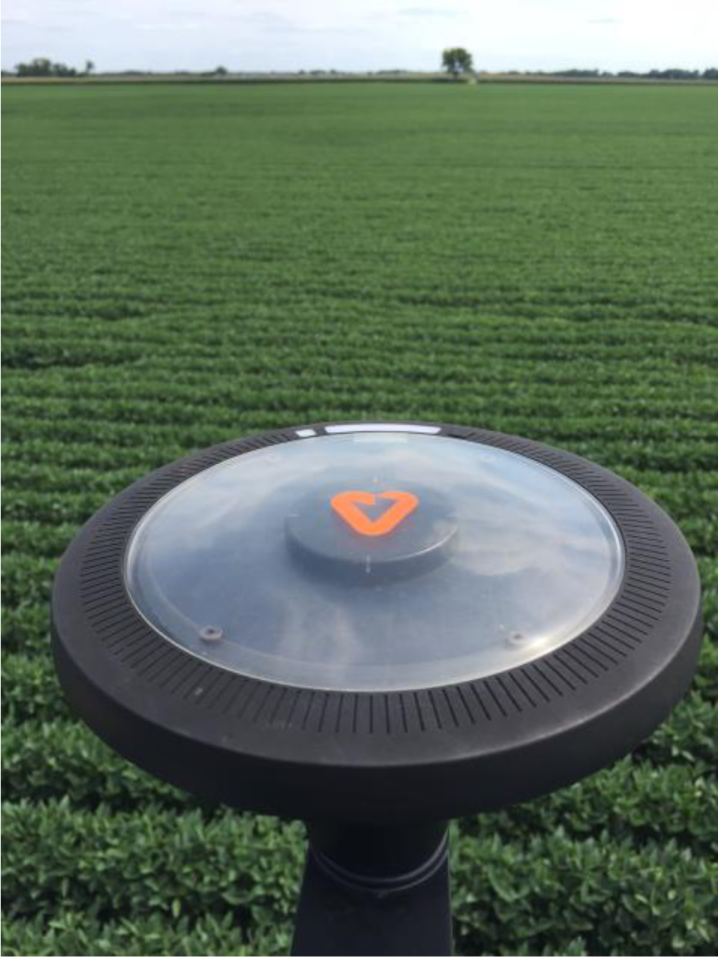
Arable Pulsepod deployed at five meter height above soybean canopy.

**Figure 2.**
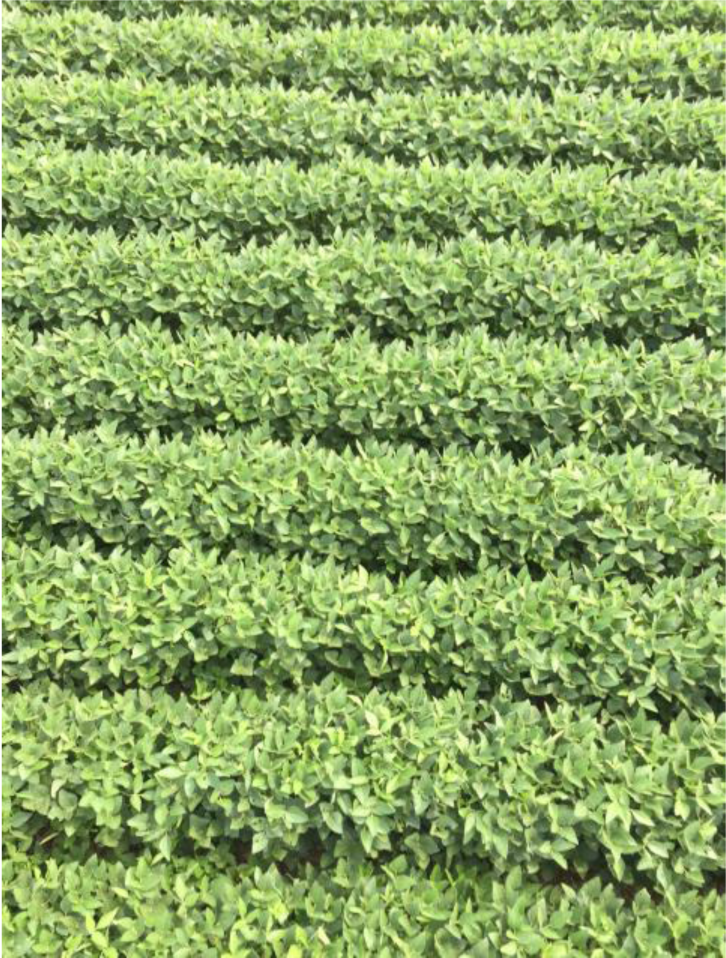
Soybean canopy below measurement location.

### Net Radiation

The components of net radiation are shortwave (downwelling and upwelling) and longwave (downwelling and upwelling). The Pulsepod measures these four components separately, using four separate sensors with upward and downward orientations. The upward facing sensors respond to incident radiation from the sun and atmosphere, and provide the downwelling measurements while the downward facing sensors respond to radiation from the Earth - the upwelling components of net radiation. The Pulsepod shortwave sensor response is calibrated against the response of a reference pyranometer, within the Kipp and Zonen CNR4 Net Radiometer (*Kipp and Zonen B.V., Delft, The Netherlands*) under field conditions. The longwave sensor responsivities have also undergone a field calibration using a reference object (sky and surface) temperature provided by the blackbody temperature equivalent of the thermal energy as measured by a reference pyrgeometer, within the Kipp and Zonen CNR4 Net Radiometer. The effective blackbody temperature is used to calculate the longwave energy using the emissivity of a blackbody (ℇ=1) and the Stefan-Boltzmann Law (8). The reference longwave measurement was compensated for changes in instrument body temperature prior to performing calibration. The longwave energy as measured by the Pulsepod is a function of a die temperature (*Tdie*) and a voltage generated in response to incident longwave energy (*Vobject*). The spectral response of the reference pyranometer is from 300 nm to 2800 nm and is calibrated according to ISO 9847:1991 and considered ISO First Class (WMO Good quality). Reference manufacturer uncertainty is stated at 5% at the daily total level (95% confidence interval) for shortwave energy and <10% at the daily total level (95% confidence interval) for longwave energy (5). Reference shortwave energy is the total radiant energy between 300 nm and 2800 nm. Reference longwave energy is the total radiant energy between 4.5 µm and 42 µm. Data shown were collected at the University of Nebraska Carbon Sequestration Program (b) Site 3 at Mead, NE in June, 2016. Device measurement height was five meters. Site coordinates are: 41.179665, -96.439336.

### Photosynthetically Active Radiation (PAR)

The Pulsepod measures downwelling PAR, which enables an understanding of the instantaneous and daily total PAR available from a source. A Pulsepod was deployed alongside an LI-190 Quantum Sensor (*LI-COR, Inc. Lincoln, NE*), with upward facing orientation. The reference sensor has a manufacturer stated accuracy of ± 5% (6). Data shown were collected at same location as net radiation measurements.

## RESULTS

Figures 3 and 4 show *Ta* as measured by a Pulsepod and by the reference instrument under the same conditions. The range of temperatures measured under the test conditions was from 16.9°C to 30.7°C. Data shown reflect a thirty-minute average of five-minute measurements (Pulsepod) and a thirty-minute average of five-second measurements (reference). The average wind speed, as reported by an anemometer mounted at the same measurement height above the canopy, varied between 0.4 m/s and 4.5 m/s over the course of the comparison with a diurnal pattern represented in Fig. 3.

**Figure 3.**
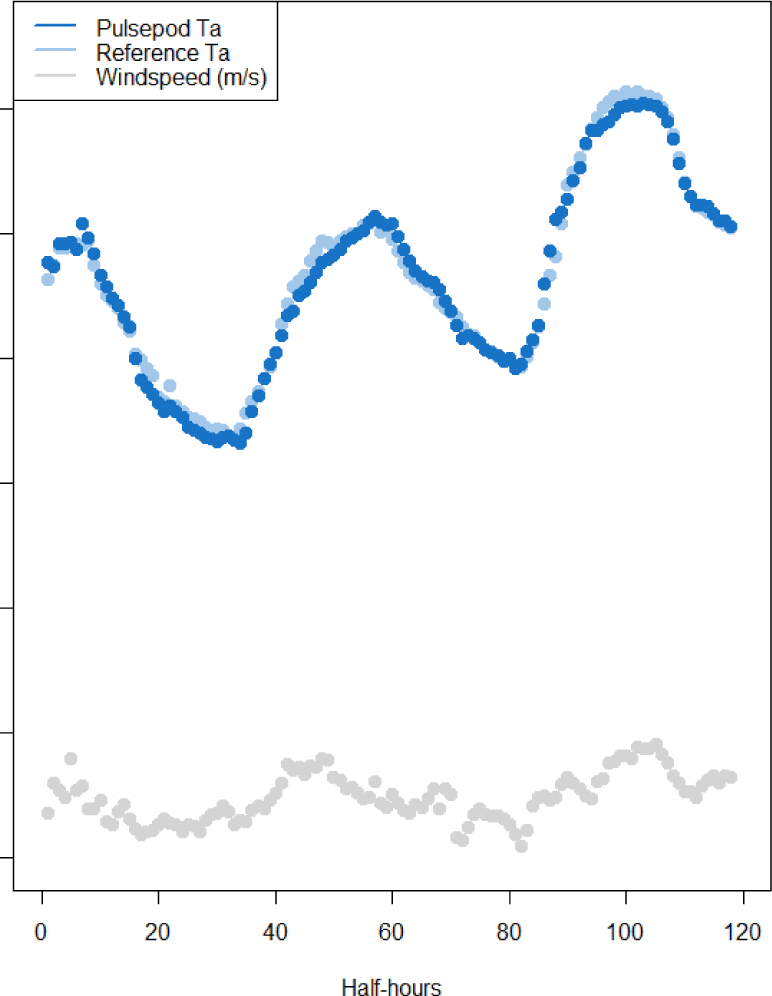
Diurnal time-series of *T*_*a*_ as measured by a Pulsepod (dark blue) and by the reference (light blue). Average windspeed is shown in gray.

**Figure 4.**
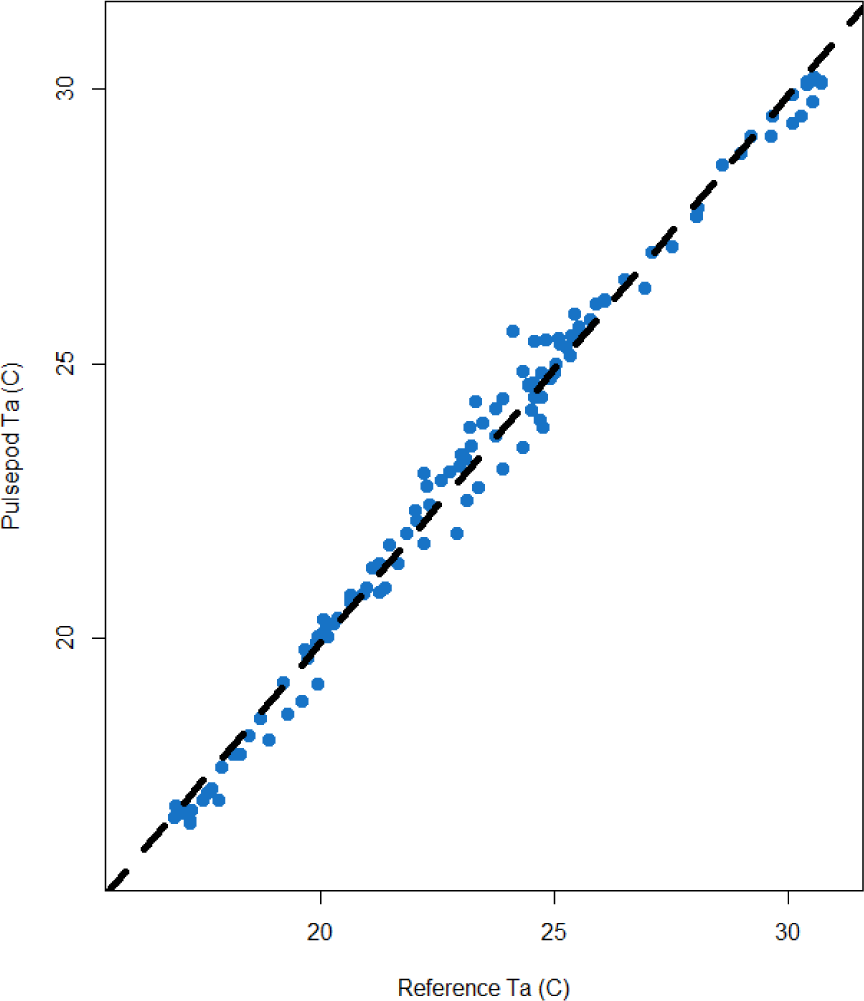
1:1 of *T*_*a*_ as measured by a Pulsepod and by the reference. R^2^: 0.99.

The relative error ratio of the Pulsepod temperature to the reference is shown in Fig. 5-6. Fig. 5 shows the ratio of measured air temperature with wind speed from 0 to 5 m/s, during periods with irradiance values less than 300 Wm^-2^. Figure 6 shows the ratio of measured air temperature during periods with wind speed from 0 to 5 m/s, with irradiance values at or above 300 Wm^-2^. During periods of time with calm conditions (< 2 m/s) and irradiance below 300 Wm-^2^, the Pulsepod reports air temperature to within +2% and -4% of the reference instrument. During periods of time with moderate wind speeds (greater than 2 m/s) and low radiative load (less than 300 Wm^-2^), the Pulsepod reports air temperature to within +3% and -2% of the reference instrument. During periods of time with moderate wind speeds (greater than 2 m/s) and greater radiative load (more than 300 Wm^-2^), the Pulsepod reports air temperature to within +6% and -4% of the reference instrument. During periods of time with calm winds (less than 2 m/s) and greater radiative load (more than 300 Wm^-2^), the Pulsepod reported a maximum relative error of +3%. The mean relative error ratio during this comparison, including all wind speed and radiation conditions, is 1.00, with a standard deviation of 0.02.

**Figure 5.**
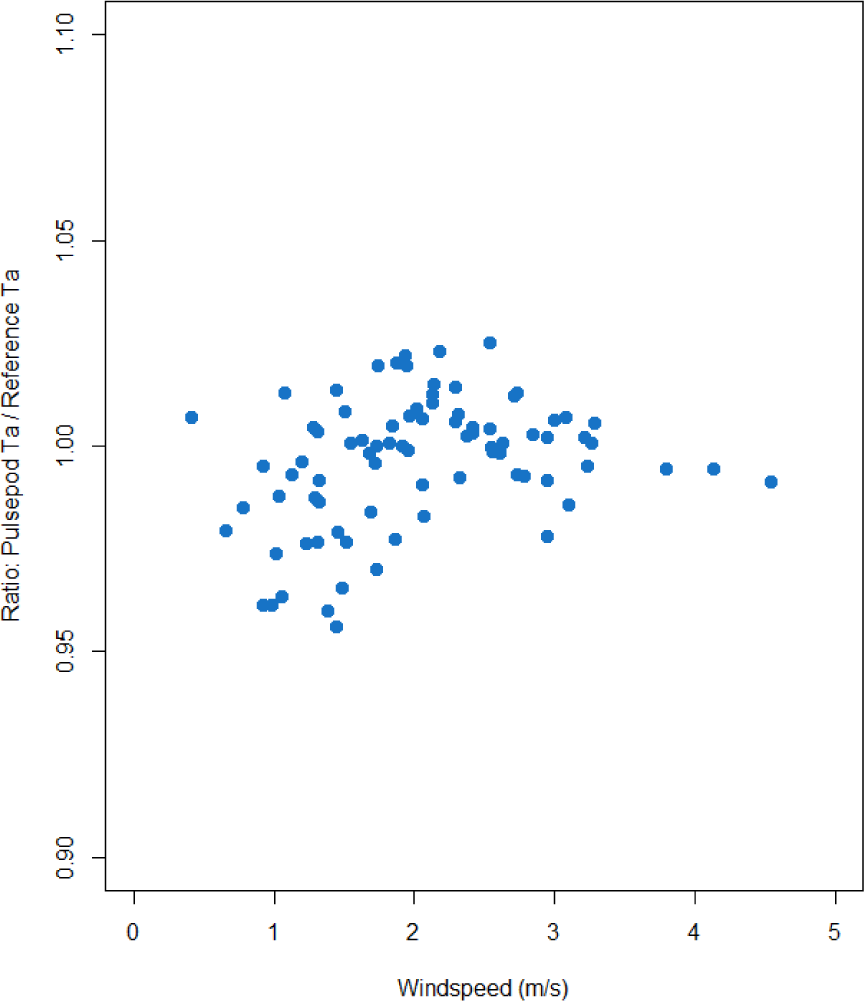
Relative error of Pulsepod *Ta* to reference *Ta* during periods with < 300 Wm^-2^.

**Figure 6.**
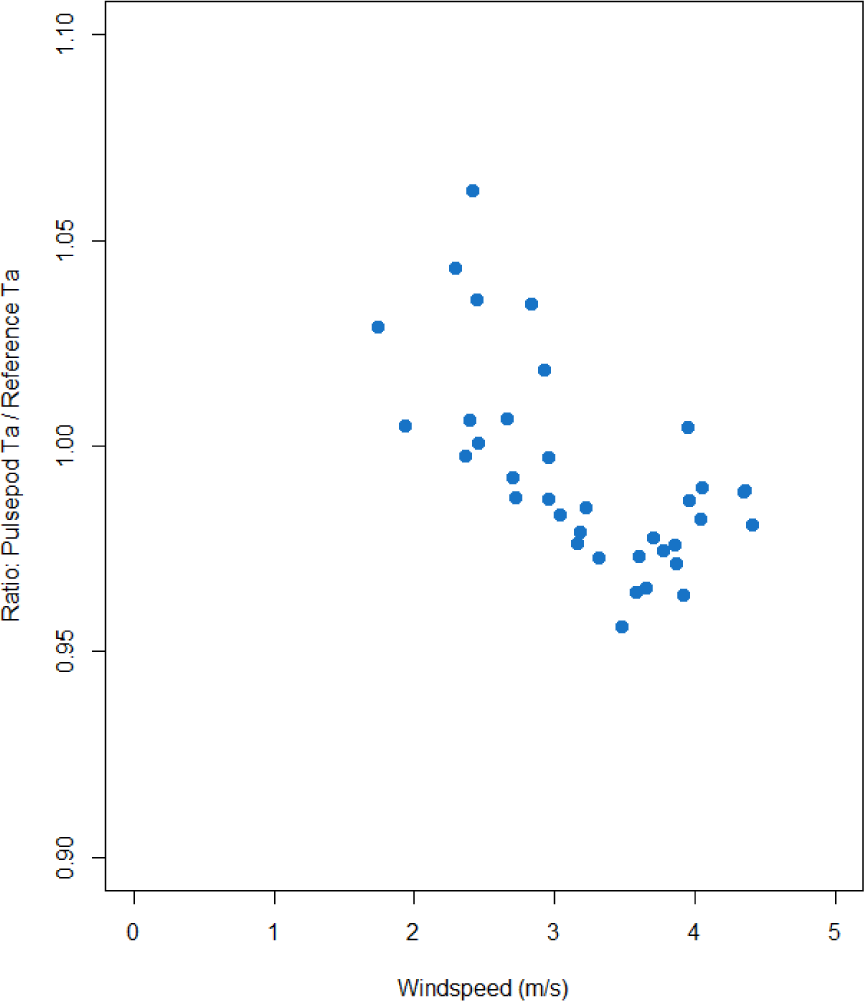
Relative error of Pulspeod *Ta* to reference *Ta* during periods with ≥ 300 Wm^-2^.

Figures 7 and 8 show *RH* as measured by a Pulsepod and the Vaisala HMP50 Temperature and Relative Humidity Sensor under the same environmental conditions. The reference has a manufacturer stated accuracy of ± 3% over the range 0 to 90% *RH* and ± 5% over the range 90 to 95% *RH* (8).

**Figure 7.**
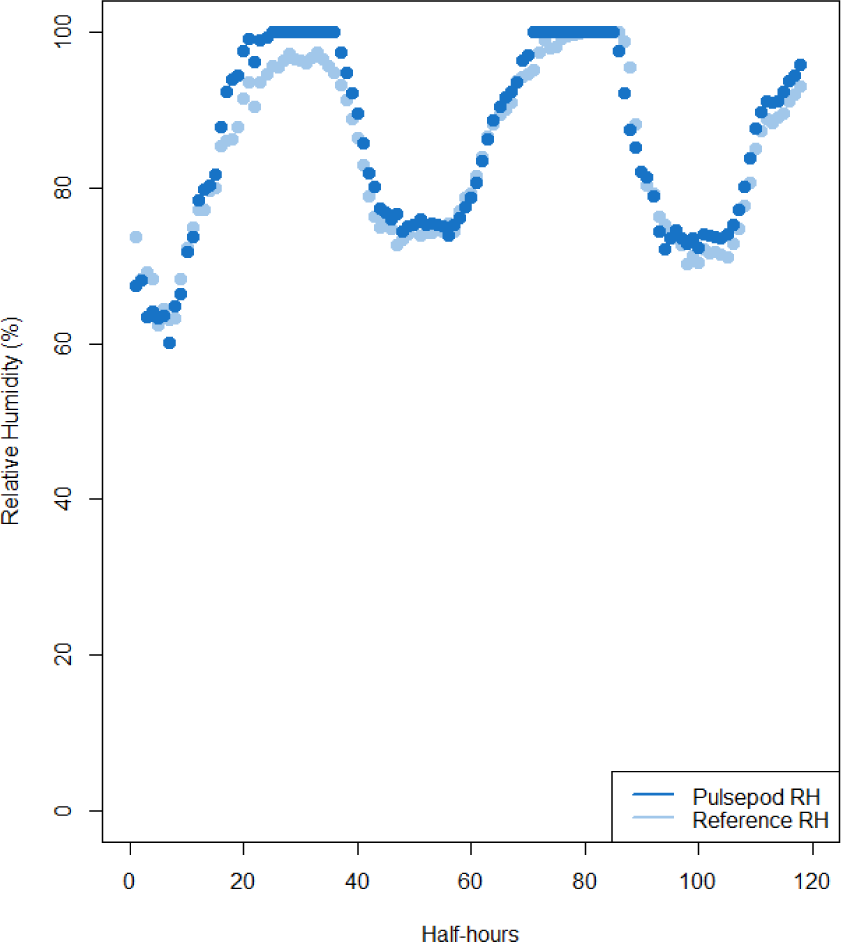
Diurnal plot of *RH* as measured by the Pulsepod and by the reference.

**Figure 8.**
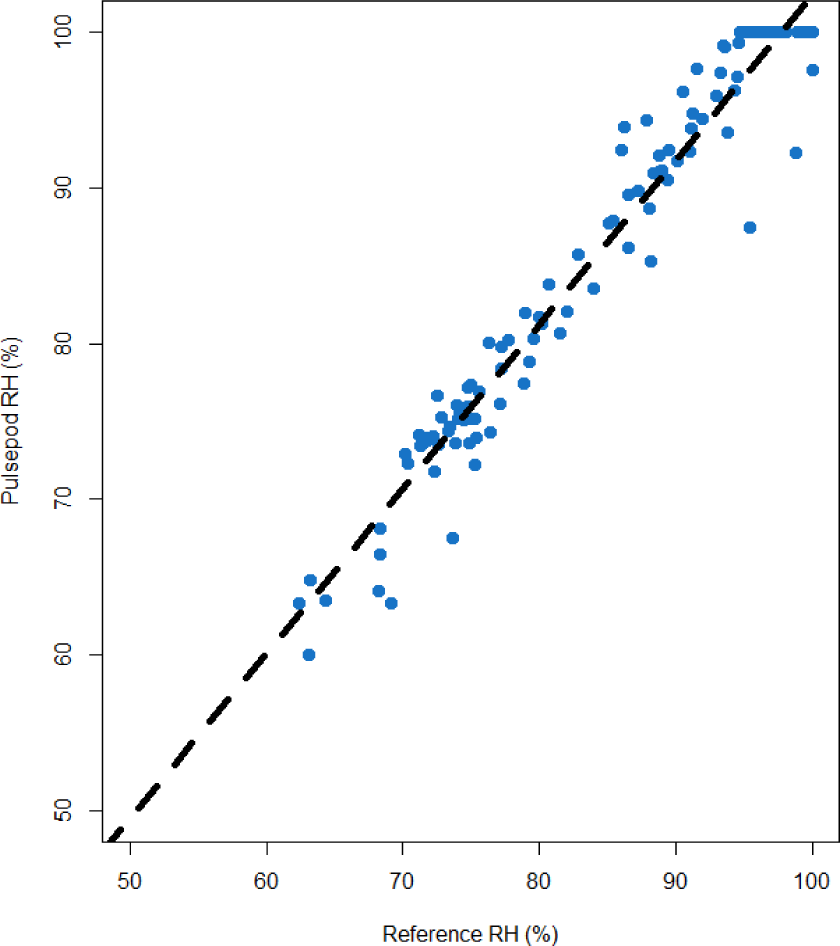
1:1 of *RH* as measured by the Pulsepod and the reference. R^2^: 0.95.

Figure 9 shows the ratio of Pulsepod *RH* to reference *RH* as a function of the ambient air temperature, and Fig. 10 shows the ratio as a function of wind speed. The maximum relative error ratio under all conditions was 1.09. The mean relative error ratio during this comparison is 1.02 with a standard deviation of 0.03. There is a reduction in relative error when wind speeds exceed 3 m/s.

**Figure 9.**
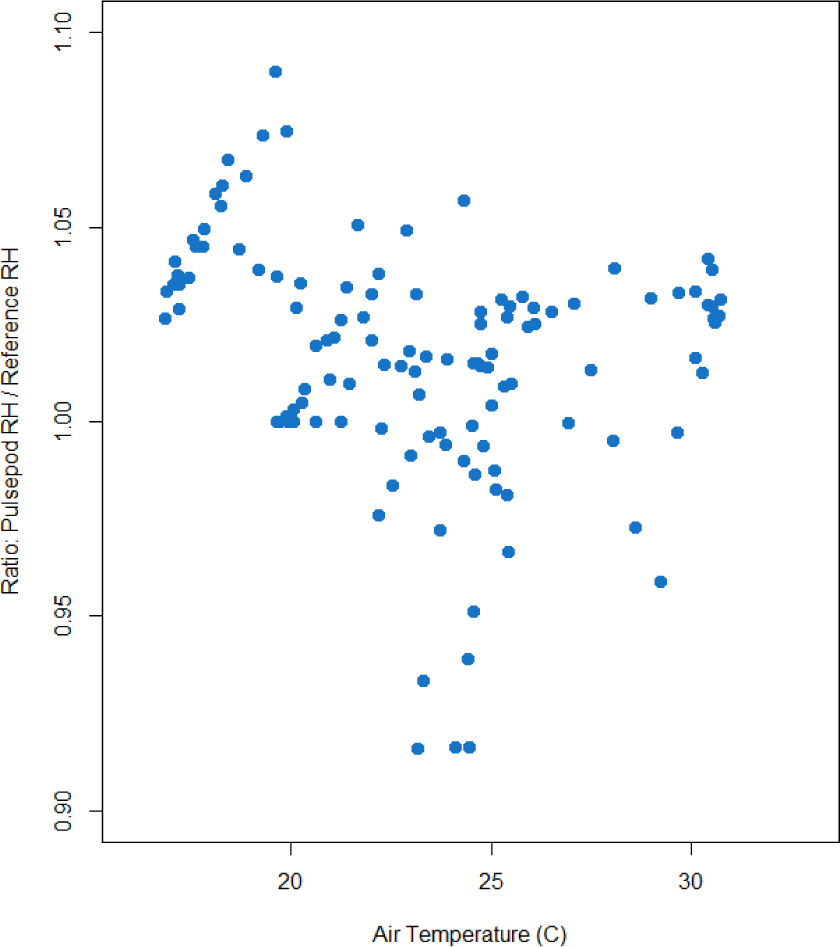
Relative error of Pulsepod *RH* to reference as a function of air temperature.

**Figure 10.**
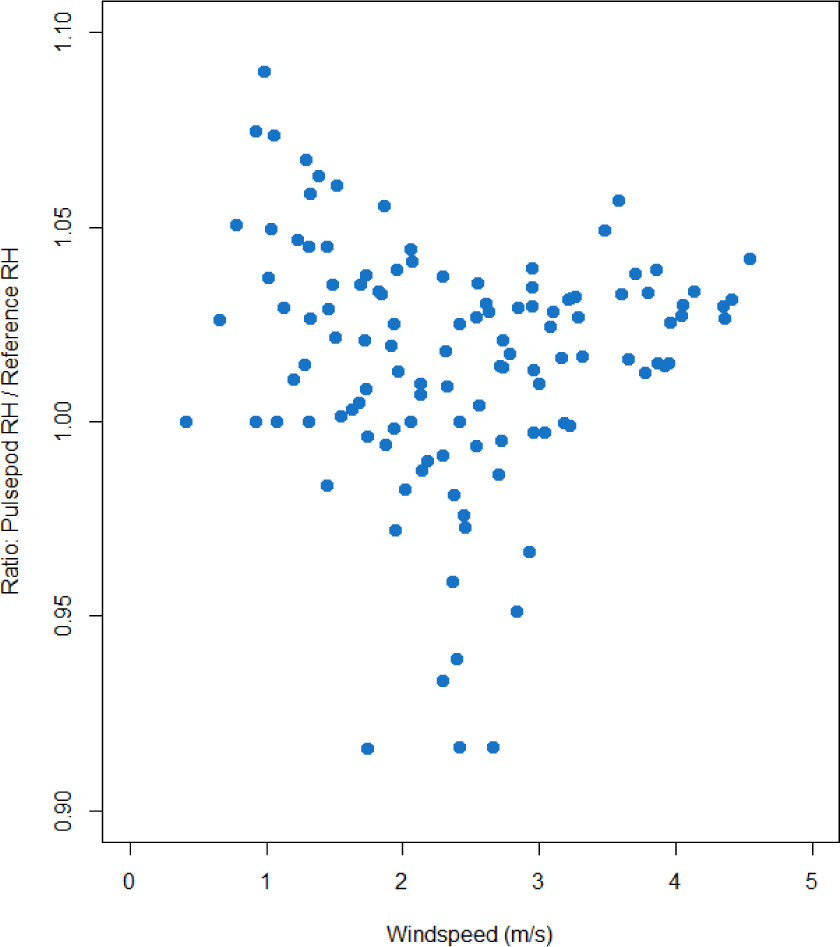
Relative error of Pulsepod *RH* to reference as a function of wind speed.

Figure 11 shows the time series of barometric pressure as measured by the Pulsepod and the reference. Figure 12 shows the 1:1 relationship between the Pulsepod and reference, and Fig. 13 shows the relative error of pressure as measured by the Pulsepod and the reference. The mean relative error ratio is 1.00 with a standard deviation of .0005.

**Figure 11.**
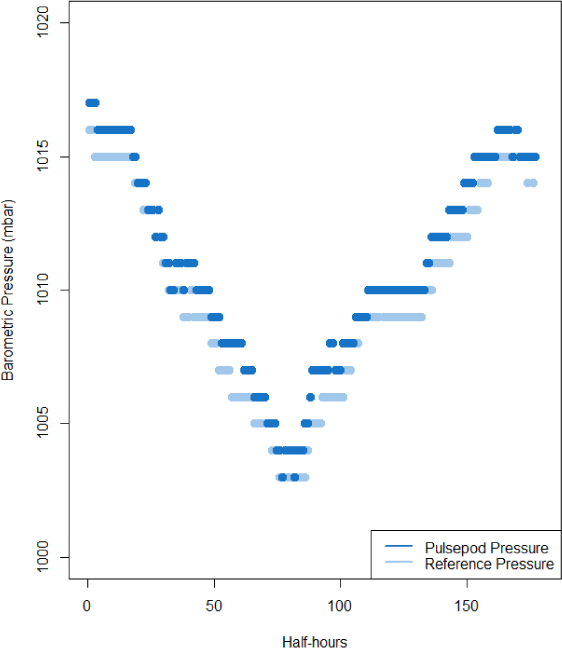
Barometric pressure as measured by the Pulsepod (dark blue) and by the reference (light blue).

**Figure 12.**
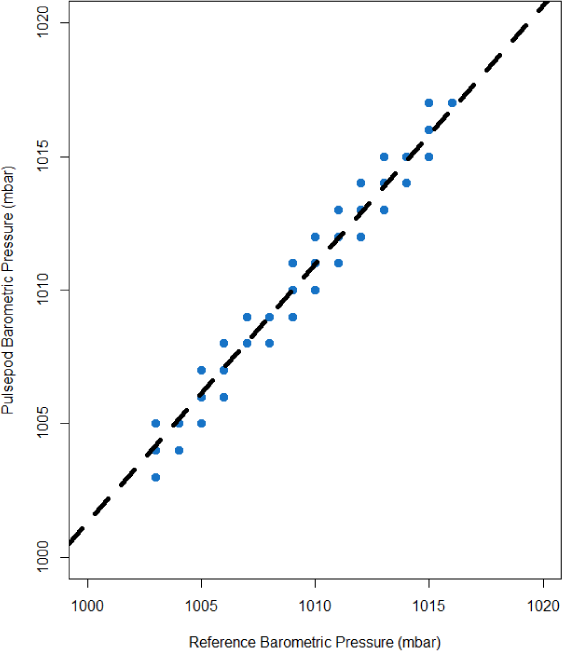
1:1 Barometric pressure as measured by the Pulsepod and reference. R^2^: 0.99.

**Figure 13.**
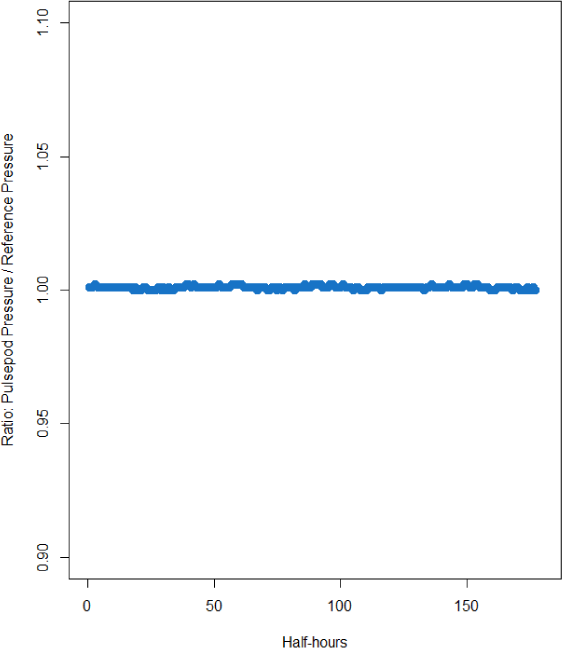
Relative error of Pulsepod barometric pressure to reference.

Figure 14 shows the time-series of longwave and shortwave energy as measured by the Pulsepod and the reference net radiometer under field conditions. Each data point represents a thirty-minute average of five-minute measurements (Pulsepod) and a thirty-minute average of five-second measurements (reference). The Pulsepod measured cumulative net radiation over the course of three separate days (Fig. 15), with a mean relative error for each day of 0.94 and standard deviation of 0.04. After 128 half-hour periods, the cumulative relative error for net radiation was also 0.94. The mean relative error at the daily level for shortwave energy (using downwelling sensors for comparison) was 0.97 with a standard deviation of 0.003. The mean relative error at the daily level for longwave energy (using the upwelling sensors for comparison) was 1.00 with a standard deviation of 0.02. Figures 16-18 show the 1:1 relationship between net shortwave energy, net longwave energy, and net radiation as measured by the Pulsepod and the reference net radiometer.

**Figure 14.**
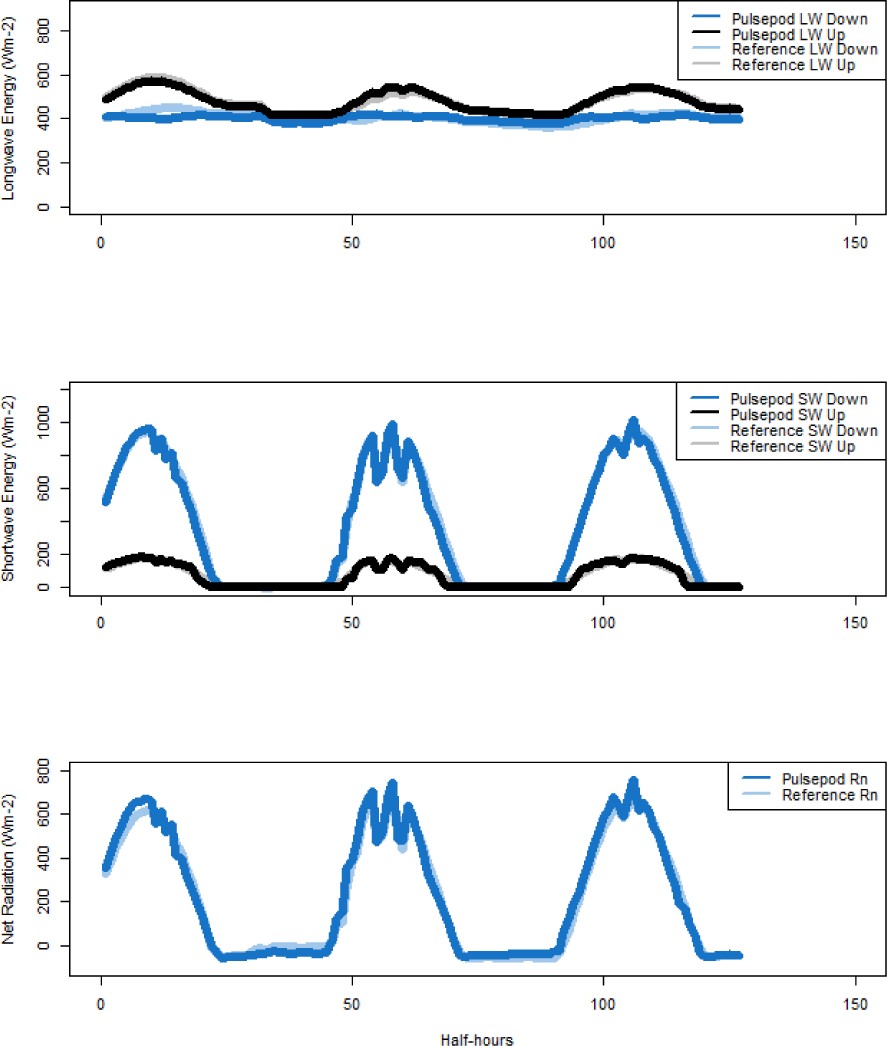
Time-series of longwave, shortwave, and net radiation as measured by the Pulsepod and the reference net radiometer under field conditions.

**Figure 15.**
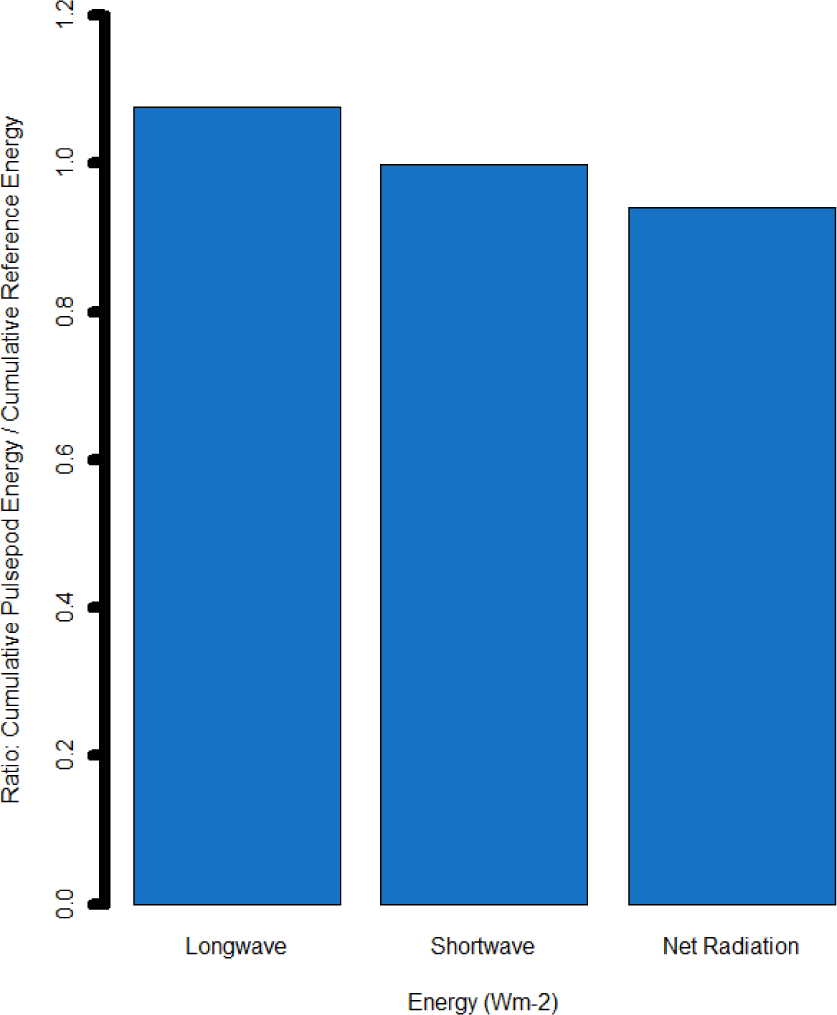
Relative error for cumulative net shortwave radiation, cumulative net longwave radiation, and cumulative net radiation over the course of three days. Cumulative relative error for net radiation after 128 half-hours was 0.94.

**Figure 16.**
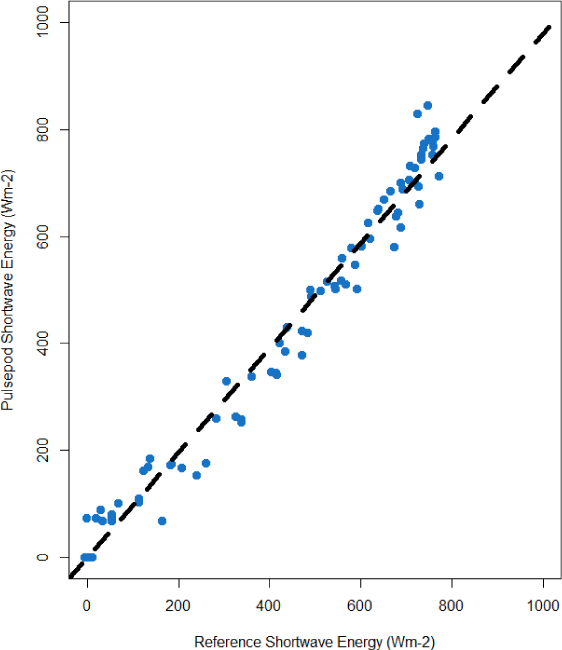
Pulsepod and reference net shortwave energy (R^2^ = 0.98).

**Figure 17.**
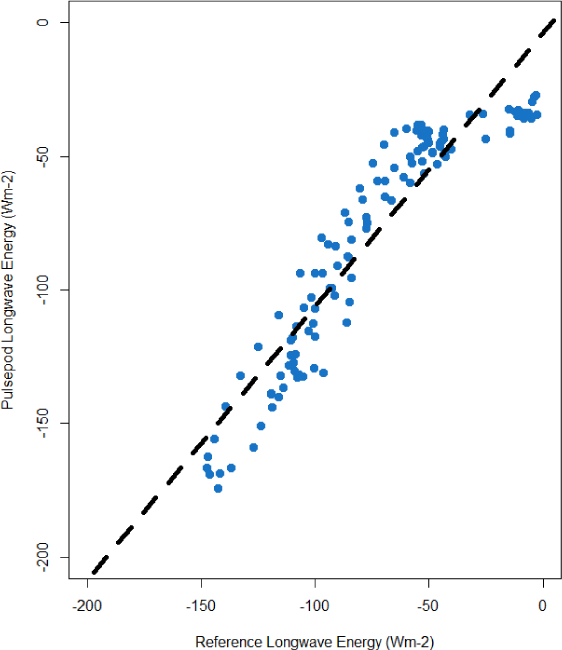
Pulsepod and reference net longwave energy (R^2^ = 0.87).

**Figure 18.**
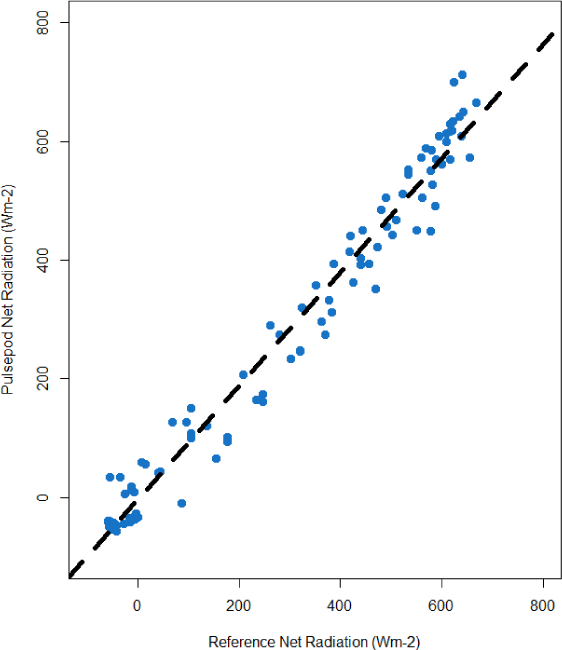
Pulsepod and reference net radiation (R^2^ = 0.98).

Figure 19 shows the timeseries of PAR as measured by the Pulsepod and the reference sensors. Figure 20 shows the 1:1 relationship between the Pulsepod and reference when measuring downwelling PAR. The Pulsepod measured cumulative downwelling PAR over the course of three separate days (Fig. 19), with a mean relative error for each day of 0.95 and standard deviation of 0.006. After 128 half-hour periods, the cumulative relative error was also 0.95.

**Figure 19.**
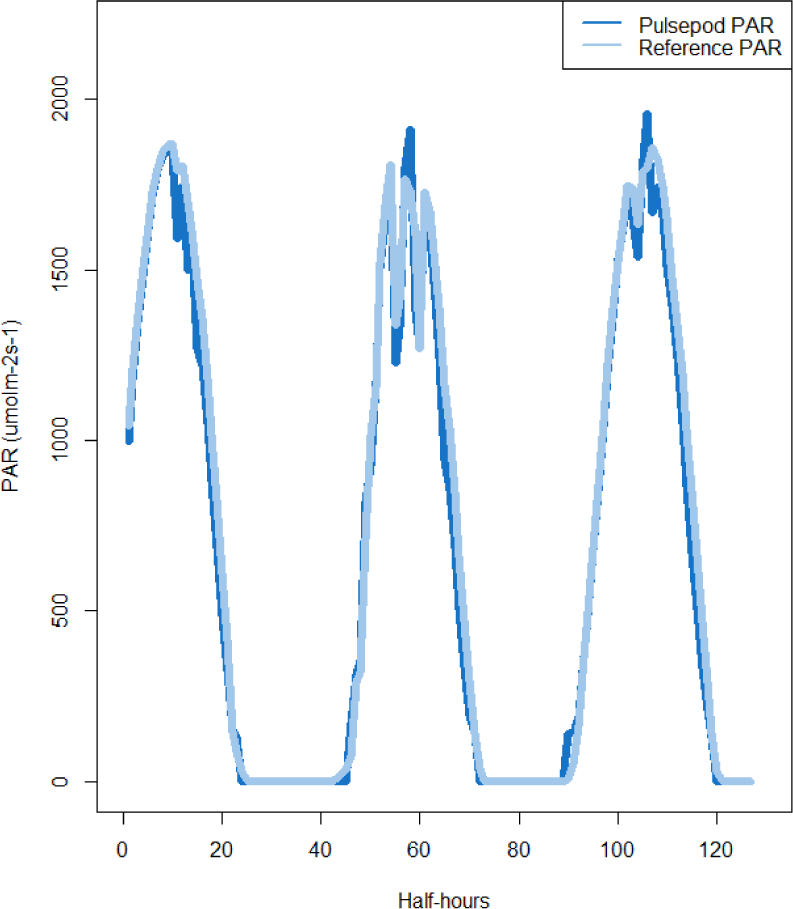
Time-series of downwelling PAR as measured by the Pulsepod and the reference sensor.

**Figure 20.**
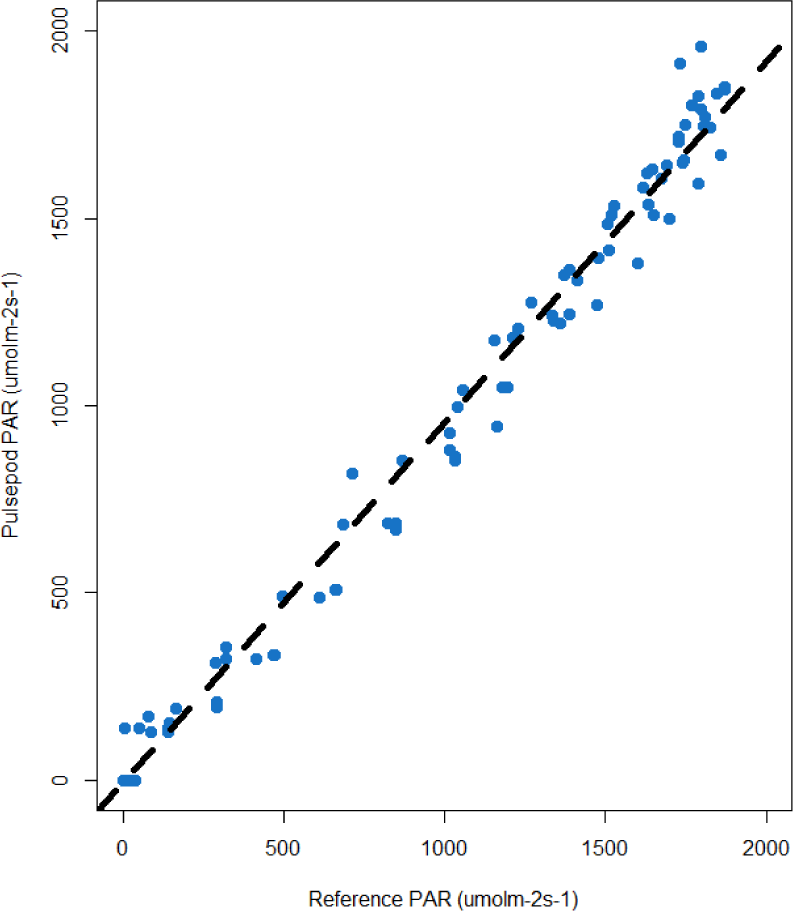
Pulsepod and reference downwelling PAR (R^2^ = 0.98)

## DISCUSSION AND CONCLUSIONS

The device reports the microclimate and radiometric measurements near or within the accuracy limits stated by the reference manufacturers for the measurements shown in this comparison. Standard weather station variables were reported to within 4% of reference (air temperature), 6% of reference (relative humidity) and 0.1% of reference (barometric pressure). The device reported net radiation to within 10% (Table 2) of the reference at the daily total level. The downwelling longwave component of radiation shows the greatest variability (Fig. 17) for net radiation components, which could be due to the changing emissivity of the sky during the course of the study. Further work could identify changing emissivity over time, and incorporate these changing values into the calculation of longwave energy for improved accuracy.

**Table 1.** Pulsepod sensor accuracy, relative accuracy, and manufacturer stated accuracy for microclimate measurements.

| Measurement | Pulsepod Sensor Accuracy | Pulsepod to Reference Relative Accuracy | Reference Manufacturer Accuracy |
| --- | --- | --- | --- |
| Air Temperature | $\pm 0.75^{\circ}\text{C}$ over the range $-40$ to $125^{\circ}\text{C}$ | $\leq 4\%$ of measurement (mean relative error ± one $\sigma$) | $\pm 1.5^{\circ}\text{C}$ over the temperature range $-40^{\circ}\text{C}$ to $60^{\circ}\text{C}$ |
| Relative Humidity | $\pm 3\%$ *RH* over the range $0$ to $100\%$ | $\leq 6\%$ of measurement (mean relative error ± one $\sigma$) | $\pm 3\%$ *RH* over the range $0$ to $90\%$ *RH* and $\pm 5\%$ over the range $90$ to $95\%$ *RH* |
| Barometric Pressure | $\pm 4$ mbar over the pressure range $500$ to $1100$ mbar | $\leq 0.1\%$ of measurement (mean relative error ± one $\sigma$) | $\leq 0.05\%$ over analog pressure range (± one $\sigma$) |

**Table 2.**
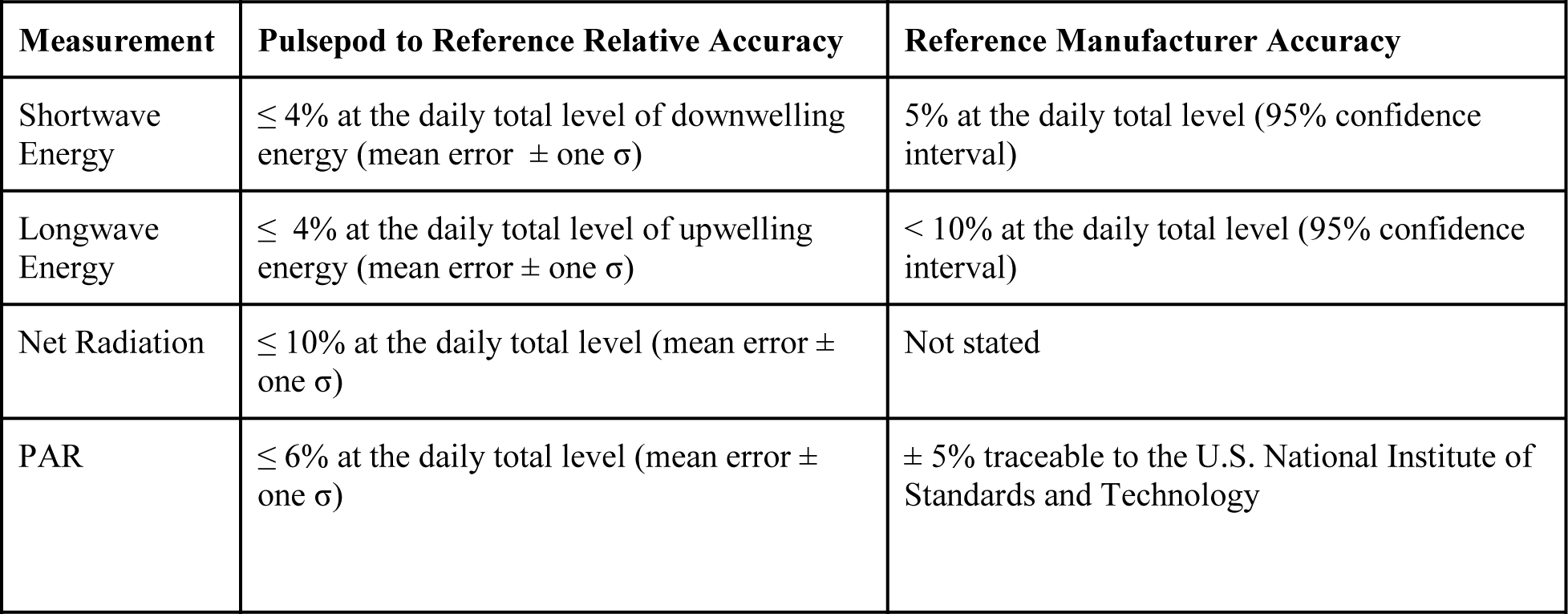
Pulsepod measurement accuracy relative to reference and manufacturer stated accuracy for radiant energy measurements.

## ACKNOWLEDGEMENTS

We gratefully acknowledge researchers at Rutgers University and the University of Nebraska in facilitating data access and field support over the course of the study.

## Footnotes

a Rutgers University Photochemical Assessment Monitoring (PAM) Site (http://pamsite.rutgers.edu/sub_data_display/met_tower.php)

b University of Nebraska Carbon Sequestration Site (UNL CSP) (http://csp.unl.edu/Public/)

